# The probiotic roles of *Lactiplantibacillus plantarum* E2 as a dietary supplement in growth promotion and disease resistance of juvenile large yellow croaker (*Larimichthys crocea*)

**DOI:** 10.1101/2023.06.24.544721

**Authors:** Liu Ruizhe, Wang Shan, Huang Dongliang, Huang Yulu, He Tianliang, Chen Xinhua

## Abstract

Bacterial diseases are the most important limiting factors for the development of large yellow croaker aquaculture. Probiotics are considered to be a promising alternative approach for the control of bacterial diseases in aquaculture. However, the studies on probiotics used in farmed large yellow croakers were very limited. In this study, we isolated and identified a *Lactiplantibacillus plantarum* E2 from the intestinal tract of large yellow croaker. *L. plantarum* E2 showed significant antibacterial activities against several aquaculture pathogenic bacteria, intestinal environmental tolerance, and biosafety. After 7 weeks of feeding, the E2 supplementation of dietary significantly improved the growth and the survival rates of large yellow croakers after *Pseudomonas plecoglossicida* PQLYC4 challenge. Further analysis showed that E2 effectively improved the intestinal integrity, and increased the intestinal α-amylase, trypsin and lipase activities. Moreover, the E2 supplementation also significantly suppressed the mRNA expression of IL-10 and increased the mRNA expression of IL-1β, IL-12α, IL- 17D, IFN-γ, and TNF-α-R. Gut microbiota analysis showed that E2 significantly affected gut microbial community composition by decreasing the relative abundance of *Sphingomonas* and increasing the relative abundance of *Lactobacillus* and *Pseudomonas.* Finally, E2 could improve resistance of large yellow croaker against *P. plecoglossicida* PQLYC4 infection. Therefore, our findings showed that *L. plantarum* E2 has potential application as a probiotic in large yellow croaker, which may provide a new strategy of preventing and controlling bacterial diseases in this species.

**Highlights:** *Lactiplantibacillus plantarum* E2 showed significant antibacterial activities against several aquaculture pathogenic bacteria, intestinal environmental tolerance, and biosafety.

*Lactiplantibacillus plantarum* E2 supplementation improved growth rates, and intestinal health of large yellow croaker.

*Lactiplantibacillus plantarum* E2 increased the abundance of potential probiotics in the intestine tract of large yellow croaker.

## Introduction

With the increasing human demand for animal protein, aquaculture, especially marine aquaculture, has become the fastest-growing food-producing industry in many countries (FAO, 2022, 2012). High density and intensification of aquaculture have led to the deterioration of the aquaculture environment, resulting in the frequent aquaculture diseases (Rodger, 2016). The economic losses caused by aquaculture diseases are estimated to reach 30 billion yuan annually in China (Xu and Lv, 2018). So far, the conventional approaches applied in aquaculture disease control mainly included the uses of antibiotics or other chemical agents (Sharma et al., 2012). However, the misuse of these chemical agents has resulted in antibiotic accumulation in aquatic products and increases of antibiotic-resistant bacteria in the environment (Cabello, 2006; Lulijwa et al., 2020). Therefore, the use of antibiotics or other chemical agents in aquaculture has been banned or severely restricted in recent years.

Probiotics are a group of living microorganisms that can benefit host health when administered in adequate amounts (Reid et al., 2003). In aquaculture, probiotics have been reported as a promising alternative approach for the control of diseases (Chauhan and Singh, 2019; Iorizzo et al., 2022). The benefits of probiotics mainly include inhibiting growth of pathogenic agents, improving the balance of gut microbiota, increasing the absorption and utilization of feed nutrients by the host, and regulating host immunity (Amenyogbe et al., 2020). Probiotics in aquaculture include lactic acid bacteria (LAB), *Bacillus*, *Carnobacterium,* and *Saccharomyces* (Abd El-Rhman et al., 2009; Abdel-Tawwab et al., 2008; El-Saadony et al., 2021; Puvanendran et al., 2021). LAB is a group of Gram-positive bacteria that produce lactic acid but do not produce spores, including many species of *Lactobacillus*, *Lactiplantibacillus*, *Bifidobacterium,* and *Enterococcus* (Iorizzo et al., 2022; Ljungh and Wadstrom, 2006). A large number of studies have provided enough evidence that LAB plays an essential role in improving the health and welfare of fish (Giri et al., 2013; Jatobá et al., 2011; Sherif et al., 2021; Hai-peng Zhang et al., 2020).

In recent years, *Lactiplantibacillus plantarum* (previously classified as *Lactobacillus plantarum*) has gained considerable attention as a probiotic used in aquaculture (Iorizzo et al., 2022). Previous studies suggested that some strains of *L. plantarum* inhibited pathogenic bacteria, regulated host gut microbiota, improved host immunity, and promoted host growth (Giri et al., 2013; Mohammadian et al., 2016; Hai-peng Zhang et al., 2020). Moreover, some *L. plantarum* strains could tolerate acid and bile conditions in the intestinal tract of fish, which were essential for their colonization and probiotic effects in the intestinal tract (Van Doan et al., 2020). Several strains of *L. plantarum,* such as *L. plantarum* C20015, VSG3, WCFS1, and CCFM8661, have probiotic effects on *Nile tilapia, Oncorhynchus mykiss, Barbus grypus, Labeo rohita* and *Cyprinus carpio* (Balcázar et al., 2008; Giri et al., 2013; Hamdan et al., 2016; Mohammadian et al., 2016; Hai-peng Zhang et al., 2020). However, these strains of *L. plantarum* may not be well adapted to all fish species due to the differences in gut environments between species. Accumulating evidences demonstrated that the host-associated probiotics were more effective than probiotics from other origins (Lazado et al., 2015; Van Doan et al., 2020). Therefore, screening probiotics for specific species of cultured fish is necessary.

Large yellow croaker (*Larimichthys crocea*) is a mariculture fish with the highest annual production and significant economic value in China (“An improved genome assembly for Larimichthys crocea reveals hepcidin gene expansion with diversified regulation and function | Communications Biology,” n.d.). The high density and intensification of the aquaculture model have led to the frequent occurrence of bacterial diseases (Li et al., 2020). However, the studies on probiotics used in farmed large yellow croakers were very limited. Ai et al. (2011) reported that administration of *B. subtilis* enhanced the growth performance and feed utilization of juvenile large yellow croaker, as well as the non-specific immunity and disease resistance. Moreover, *Clostridium butyricu*m as a probiotic could improve intestinal development, immune response, and gut microbiota in large yellow croaker larvae (Yin et al., 2021). However, the LAB, especially host-associated LAB in large yellow croaker, has not been reported. In this study, we obtained an *L. plantarum* E2 from the intestinal tract of large yellow croaker. The antibacterial activity, intestinal environmental tolerance, biosafety, growth promotion, and visceral white nodules disease resistance of *L. plantarum* E2 have been evaluated. Furthermore, we also analyzed the change of large yellow croaker gut microbiota after *L. plantarum* E2 feeding. Our findings may provide a new method for preventing and controlling bacterial diseases in large yellow croaker.

## Materials and methods

### Isolation of intestinal microorganisms from large yellow croaker

In July 2022, the farmed large yellow croakers were collected at marine net cages in Ningde, China. Healthy fish with a body weight of 100-500 g were euthanized by immersion in a tank containing 150 mg/L MS-222 (Sigma, USA) for 15 min. Then, the intestinal tract of fish was aseptically collected into a sterile sample tube, and stored in ice. After being transported to the laboratory, the intestinal sample was homogenized in 2 mL of 10 mM phosphate-buffered saline (PBS, pH 7.2) solution for 2 min at room temperature. Homogenates were spread over the surface of triplicate plates of LB agar, 2216E agar, tryptic soy agar (TSA), and De Man Rogosa and Sharpe agar (MRS) with incubation at 28℃ for 3–7 days. Several colonies were observed and isolated randomly on fresh media based on their morphology. The pure cultures were obtained by streak plate isolation and stored at −80℃ in the corresponding isolation medium contained 20% (v/v) glycerol. All culture media were purchased from Haibo Biotechnology Company (Qingdao, China).

The animal study in the present experimental was reviewed and approved by Animal Care Committee, Fujian Agriculture and Forestry University (Fuzhou, Fujian, China, approval ID 202003016).

### Tested strains

Five important bacterial pathogens from large yellow croaker, including *Vibrio alginolyticus* (MCCC 1A03220), *Vibrio harveyi* (MCCC 1A03227), *Vibrio campbellii* (MCCC 1A08161), and *Aeromonas hydrophila* (MCCC 1A00007) were obtained from the Marine Culture Collection of China. Moreover, *Pseudomonas plecoglossicida* strain PQLYC4, which was isolated and preserved in our lab (Li et al., 2020), was also used to test antimicrobial activity of the isolated intestinal strains. *P. plecoglossicida* and *A. hydrophila* were cultured in tryptic soy broth (TSB, Haibo Biotechnology Company) medium at 28℃, while *V. alginolyticus*, *V. harveyi*, and *V. campbellii* were cultured in Zobell 2216E (2216E, Haibo Biotechnology Company) medium at 28℃.

### Antimicrobial activity detection

The agar well diffusion method was performed to detect the antimicrobial activity of the isolated intestinal microorganisms according to our previous study (Wu et al., 2018). Briefly, each tested strain was cultured to the logarithmic growth phase. When 100 mL of 2216E, TSB or MRS with 0.75% (w/v) agar was sterilized and cooled down to 45℃, 100 μL of the tested strain culture was added and mixed by shaking slightly. The mixture was poured into sterile plates. After solidification, a well with an 8 mm diameter was made in center of each agar plate and then was added 100 µL of the culture supernatant of each isolated intestinal strain. The plates were cultured at 28℃ for 72 h to observe for antimicrobial activity. Each test was repeated in triplicate.

### Identification of strain E2

Genomic DNA of the strain E2 was extracted using a TIANamp Bacterial DNA Kit according to the manufacturer’s protocol (Tiangen Biotech Co., LTD, Beijing, China). The PacBio and Illumina Miseq sequencing of the strain E2 genome and bioinformatic analysis were performed by Guangdong Magigene Biotechnology Co., Ltd. (Guangzhou, China). For homology analysis, the 16S rDNA sequence of the strain E2 was aligned with NCBI rRNA/ITS databases using the blastn algorithm (http://blast.ncbi.nlm.nih.gov/blast.cgi). For phylogenetic analysis, the 16S rDNA sequence of the strain E2 was aligned with that of 12 *Lactiplantibacillus* strains using the MAFFT program with default parameters. Phylogenetic tree was constructed with maximum likelihood estimate method (1,000 bootstrap replicates) using the MEGA 11 software. Average nucleotide identity analysis based on MUMmer (ANIm) was performed between the genomes of the strain E2 and other 12 *Lactiplantibacillus* strains using Jspecies (version 1.2.1).

### Hemolytic assay

To confirm the potential pathogenicity of the strain E2, a hemolysis assay was performed as previously described (2016). In brief, overnight culture of the strain E2 was streaked on 7% sheep blood agar plates (Haibo Biotechnology Company). These plates were then incubated anaerobically at 37℃ for 24 h and examined for signs of hemolysis. *Staphylococcus aureus* (ATCC 25923) was used as positive control.

### Sensitivity to pH and gastrointestinal tract conditions

The strain E2 was grown to log phase in MRS medium with 1.5% NaCl at 28℃ before the assay. For resistance to pH, strain E2 was cultured in MRS medium adjusted to three pH levels (pH 3.0, 7.0, and 9.0) using 0.1 N HCl and 1 M NaOH. For resistance to gastrointestinal tract conditions, the simulated gastric juice (SGJ) (3 mg/mL pepsin adjusted to pH 3.0) and simulated intestinal juice (SIJ) (1 mg/mL pancreatin, 1 mg/mL bile salts and adjusted to pH 8.0) was evaluated following the modified methods of Ratchanu et al (2021). The survived bacterial cells of each LAB were enumerated by dilution plate count on MRS agar after exposure to SGJ and SIJ, respectively. After cultured at 28℃ for 24 h, the sensitivity to pH and bile salts were determined by assessing bacterial growth at 620 nm using a V-T3 spectrophotometer (YiPu Instrument Manufacturing Co., LTD, Shanghai, China). Ability to survive was determined by comparing with the initial cell concentration at 0 h. The percentage survival was calculated according to the following equation: Survival (%) = (log_10_ CFU ml^−1^ of viable cells before treatment ÷ log_10_ CFU ml^−1^ of viable cells after treatment) × 100.

### Antibiotic sensitivity test

A total of 11 commercially available antibiotic discs (Hangzhou Microbial Reagent Co., LTD, Hangzhou, China) including Tetracycline (30 μg/disc), Clindamycin (2 μg/disc), Penicillin (10 μg/disc), Cefalexin (30 μg/disc), Erythromycin (15 μg/disc), Chloramphenicol (30 μg/disc), Amoxicillin (20 μg/disc), Vancomycin (30 μg/disc), Enrofloxacin (10 μg/disc), Kanamycin (30 μg/disc), and Norfloxacin (10 μg/disc) were used to test the antibiotic sensitivity of strain E2. Strain E2 was cultured to the logarithmic growth phase, the MRS with 0.75% (w/v) agar was mixed with the strain E2 culture as previously described. The overnight culture of strain E2 was interpreted by the disc diffusion method on agar plates according to the method described in the previous study (Sherif et al., 2021). Each test was repeated in triplicate.

### Diet Formulation

According to our previous study (Zhu et al., 2021), the basic diet containing approximately 45% crude protein and 11% lipid was formulated as the control. The culture of *L. plantarum* strain E2 (approximate 1.35 × 10^10^ CFU/ml) was supplemented to the basal diet to obtain the E2 diet (containing 1.35 × 10^9^ CFU/g) (**Supplementary Table 1**). The diet preparation process was carried out as described by Zhu et al (2021). Chemical analysis of experimental feed was performed according to the standard methods.

### Feeding trial

A total of 222 healthy large yellow croaker juveniles were obtained from the Fujian Yangze Marine Biotechnology Co. Ltd (Fuzhou, China) and reared in the culture ponds at the company. Before the start of the experiment, the fish were fed with control diet for 2 weeks at a rectangular concrete pond (4.0 × 8.0 × 1.5 m).

At the initiation of the experiment, fish with a weight of 3.15 ± 0.07 g and 3.20 ± 0.07 g in were randomly divided into 2 groups (control group and E2 group). Each group comprised 3 concrete ponds with 37 fish per pond. During the experimental period, all ponds kept uninterrupted aeration, with a water temperature of 25-31℃, a salinity of 25-27, and a dissolved oxygen of approximately 6 mg/L. The water volume was renewed twice daily. Fish were hand-fed to apparent satiation twice daily (06:00 and 18:00). The feeding trial lasted for 7 weeks.

### Sample collection

Before the experiment was conducted, initial body weight and body length of 100 randomly collected fish were measured. At the end of the experiment, fish were fasted for 24 h before sampling. Survival rate was calculated by comparing the number of remaining fish in each pond with the initial number. Final body weight and final body length of 100 fish randomly collected from each pond were measured. 6 fish were collected from each pond and preserved in 4% paraformaldehyde for hematoxylin and eosin (H&E) staining. 36 fish from each pond were dissected on ice to obtain a crude mixture of intestines for real-time PCR and enzyme activity assays. The whole intestine of 36 fish from each pond was separated aseptically under a dissecting microscope for the analysis of digestive enzyme activity, intestinal microflora and gene expression assays.

### Histology analysis of intestine

We collected intestinal tissue samples from large yellow croakers and promptly preserved them in 4% paraformaldehyde for 24 hours. Subsequently, the samples were dehydrated and embedded in paraffin wax for further analysis. The sections with a 4- μm thickness were cut by a sliding microtome (Thermo Fisher Scientific, USA) and stained with H&E as previously reported (Zhu et al., 2021). The stained sections were examined using a Nikon Eclipse Ci light microscope with a Nikon digital sight DS-FI2 camera (Nikon).

### Expression analysis of immune-related genes

To analyze the expression change of the immune related genes (IL-1β, IL-12α, IL- 17D, IFN-γ, TNF-α-R, and IL-10) in intestinal tissues of large yellow croakers fed with the control diet and E2 diet, real-time PCR was performed by the SYBR green method. Total RNA was extracted from the pooled intestine tissues of 6 fish in each pond above using TRIzol Regent (Thermo Fisher, USA). Primers were designed based on the coding sequences of these genes from large yellow croaker genome database (Ao et al., 2015) (**Supplementary Table 2**). Real-time PCR was conducted by a real-time PCR system (QuantStudio 5, Thermo Fisher, USA) with reaction conditions as follows: 95℃ 30 s, 40 cycles of 95℃ 6 s and 60℃ 25 s. After obtaining real-time PCR data, the 2^−ΔΔCt^ method was employed to calculate gene expression level (Ding et al., 2016). All PCR reactions were performed using three biological replicates and each biological replicate included three technical replicates.

### Analysis of biochemical composition

The visceral mass was weighed and homogenized in PBS solution (pH 7.2). The proportion of tissue (g) and saline (ml) was 1:9. The homogenate of fish was then centrifuged at 4,000g for 15 min and the supernatant was collected for the following assays (Zhu et al., 2021). Trypsin activity was assayed with Trypsin Assay Kit (Nanjing Jiancheng Bio-Engineering Institute, China). The α-Amylase activity was assayed with α-Amylase Assay Kit (Nanjing Jiancheng Bio-Engineering Institute, China). Lipase activity was detected with Lipase Assay Kit (Nanjing Jiancheng Bio-Engineering Institute, China). Protein was determined by the Total protein quantitative assay kit (Nanjing Jiancheng BioEngineering Institute, China).

### Diversity analysis of intestinal bacterial community

Total bacterial DNA of the intestine samples with retained contents was extracted using an E.Z.N.A. ®Stool DNA Kit (Omega, Norcross, GA, USA). After measurement of the concentration and quality of the extracted DNA using a Nanodrop 2000c spectrophotometer (Thermo Fisher, Waltham, MA, USA), the V4-V5 region of the bacterial 16S rDNA gene was amplified via the PCR method using the primers of 515F (5′GTGCCAGCMGCCGCGGTAA-3′) and 806R (5′-CCGTCAATTCCTTTGAGTT T-3′). The high throughput sequencing for the qualified amplicon was performed on the Illumina NovaSeq 6000 platform at Guangdong Magigene Biotechnology Co., Ltd. (Guangzhou, China). Complete data were submitted to the NCBI Sequence Read Archive (SRA) database under accession number PRJNA949858. Paired-end reads were assigned to samples based on a unique barcode and truncated by cutting off the barcode and primer sequence. The raw tags were then produced via FLASH (V1.2.7) (Magoč and Salzberg, 2011). Sequences were analyzed with the UCHIME algorithm (Edgar et al., 2011) and QIIME (Caporaso et al., 2010). The effective tags were filtered and clustered into operational taxonomic units (OTUs) under a 97% nucleotide similarity level. The taxonomic annotation of OTUs was performed using Uparse software (Edgar, 2013). Venn diagram was constructed to describe the core components of the genera. Linear discriminant analysis effect size (LEfSe) was used to identify significant differences in the relative abundance of bacterial taxa (Chang et al., 2022).

### *In vivo* challenge experiments with *P. plecoglossicida*

Thirty large yellow croakers were randomly selected from each pond in the control diet and E2 diet groups. All 180 fish were acclimatized in separate tanks with aerated sea water at 24-26°C for 2 days. Before the challenge, *P. plecoglossicida* strain PQLYC4 was cultured in TSB for 12 h. Then the bacteria were collected, washed twice and then resuspended in sterile PBS to a concentration of 3.0 × 10^8^ CFU/mL. Fish were exposed to water supplemented with 1 × 10^5^ CFU/mL of strain PQLYC4 for 1.5 hours. Then, the water in tanks were completely renewed to remove the strain PQLYC4. Fish were fed with basal diets for 6 days, and the mortality of the fish was recorded for 6 days after exposure.

## Results

### Identification of the strain E2 with antibacterial activity

An antibacterial strain E2 was isolated from the intestinal tract of the farmed large yellow croakers in Ningde, China. The fermentation supernatants of the strain E2 showed obvious inhibitory activity against five aquaculture pathogenic bacteria, including *P. plecoglossicida, V. campbellii, V. harveyi, V. alginolyticus,* and *A. hydrophila* (**Figure 1A**). The mean diameters of the inhibition zone ranged from 20.6 to 28.7 mm (**Supplementary Table 3**). After the complete genome sequencing, the 16S rDNA sequence of the strain E2 (accession no. CP110242, locus_tag = OLJ37_02210) showed the identity of 99.93% and 99.87% with that of *L. plantarum* strain NBRC 15891 (NR_112690.1) and *L. plantarum* strain JCM 1149 (NR_115605.1) (**Supplementary Table 4**). The phylogenetic tree based on the 16S rDNA sequences also showed that the strain E2 clustered into one branch with *L. plantarum* strain JCM 1149 and *L. plantarum* NBRC 15891 (**Figure 1B**).

**Figure 1.**
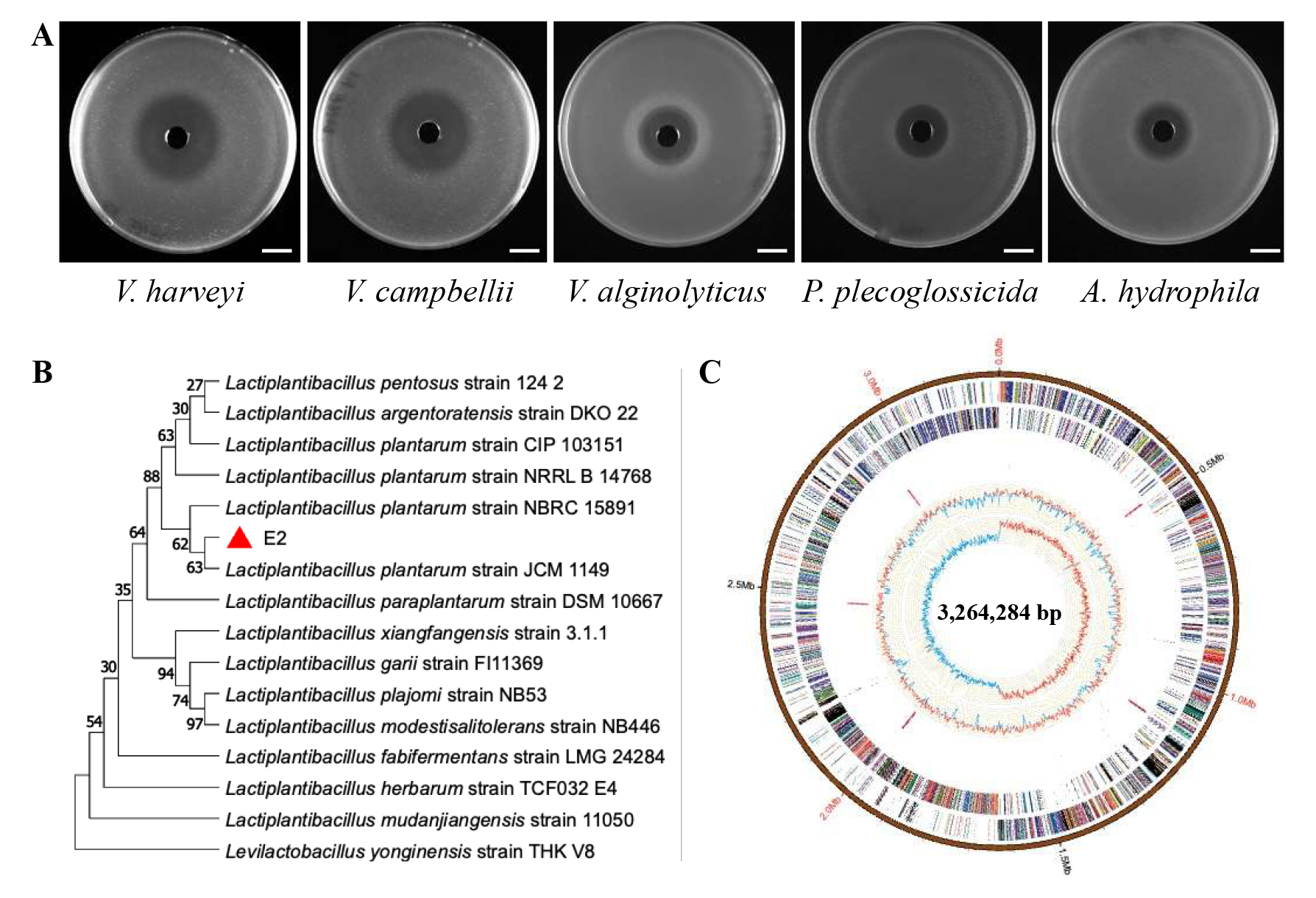
Antimicrobial activity and taxonomy identity of the strain E2. **(A)** The antibacterial activity of strain E2 fermentation supernatants against the five aquaculture pathogenic bacteria. **(B)** Phylogenetic tree of the strain E2 based on the 16S rRNA gene sequences. The tree is constructed with the maximum likelihood estimate method by the MEGA 11 software. The numbers on the nodules represent frequency of 1,000 bootstrap replicates. **(C)** Circular representation of the strain E2 genome. From outside to inside, the first circle indicated the coordinates in base pairs (bp). The second and third circles showed the predicted coding regions on positive and negative strands, respectively. The coding sequences were labelled with different colors according to their COG functions. The fourth circle showed the tRNA and rRNA genes. The fifth and sixth circles showed the GC content plot and the GC-skew.

These results suggested that the strain E2 may be a member of genus *Lactiplantibacillus*.

To further characterize the taxonomy of the strain E2, the strain E2 genome (accession no. CP110242, **Figure 1C**) was used to perform the analysis of average nucleotide identity based on MUMmer (ANIm). The ANIm value between the strain E2 and *L. plantarum* strain ST-III (NC_014554.1) was up to 99.31% (**Supplementary Table 5**), strongly suggesting that the strain E2 isolated here should be a member of *L. plantarum*, tentatively named *L. plantarum* E2.

### Biological characteristics of *L. plantarum* E2

The results of bacterial growth curve showed that *L. plantarum* E2 grew faster at 37℃ and 28℃, reaching the plateau phase for 14 h and 24 h, respectively, after inoculation. However, the strain E2 could also keep growth at 16℃. After incubation for 30 h, the bacterial content increased more than 3 times (**Figure 2A**). The strain E2 could grow at pH 3.0-7.0, yet its growth was inhibited at pH 9.0 (**Figure 2B**). The optimal pH condition was found to be at pH 5.0 and pH 7.0. The strain E2 could also grow in the simulated gastric fluid (3 mg/mL pepsin, pH 3.0) and the simulated intestinal fluid (1 mg/mL pancreatin, 1 mg/mL bile salts, pH 8.0), suggesting that strain E2 could tolerate gastrointestinal tract conditions (**Figure 2C**).

**Figure 2.**
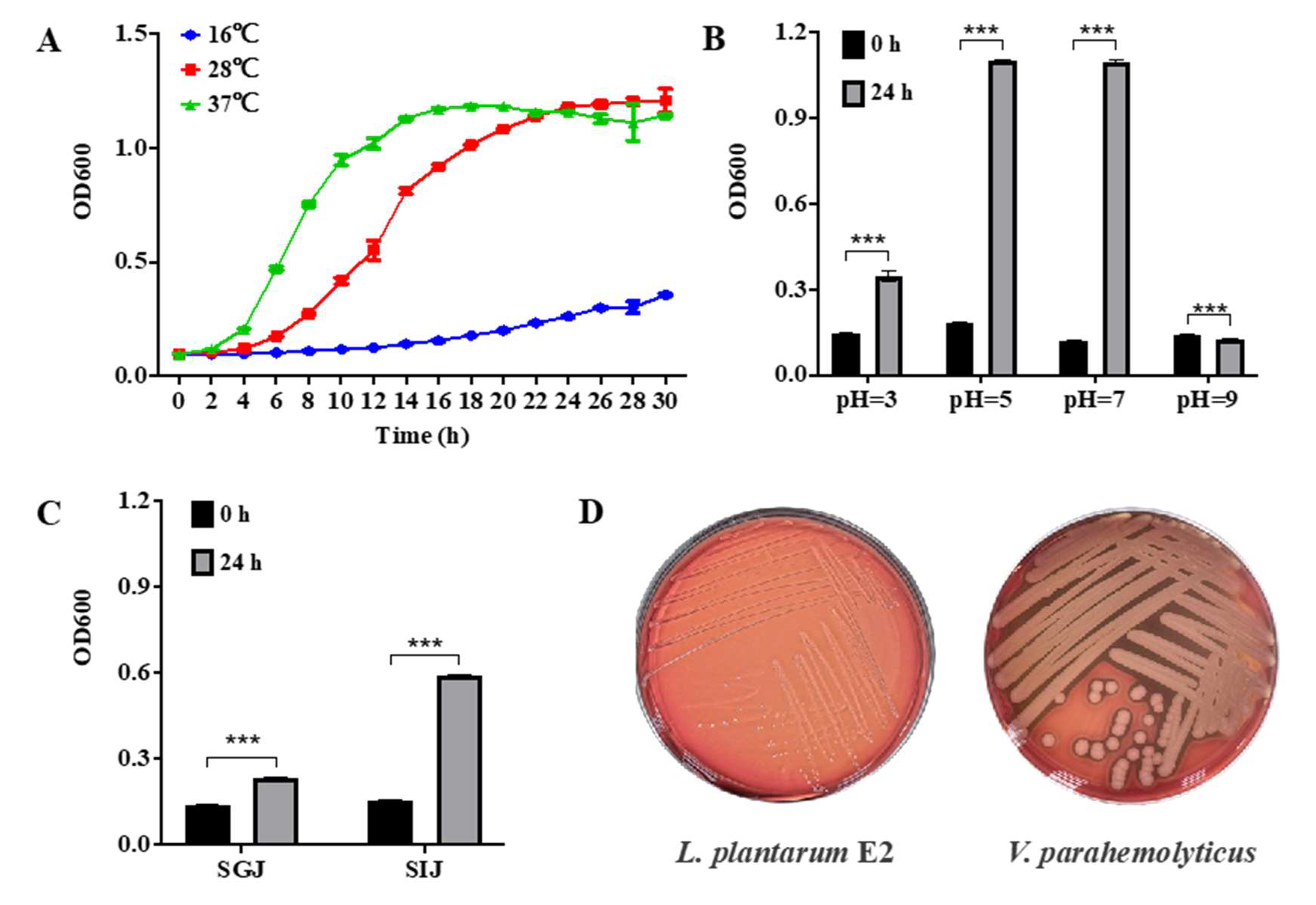
Biological characteristics of *L. plantarum* E2. **(A)** One-step growth curves of *L. plantarum* E2 at three different temperatures. **(B)** Final optical density (OD600 nm) reached by *L. plantarum* E2 grown for 24 h at 30℃in MRS broth with different values of pH (3.0, 5.0, 7.0, and 9.0). **(C)** Final optical density (OD600 nm) reached by *L. plantarum* E2 grown for 24 h at 30°C in MRS broth supplemented with 0.1% of bile salts. **(D)** Hemolytic activity of *L. plantarum* E2 compared with *V. parahaemolyticus*.

The results of hemolytic assay showed that the strain E2 did not induce any hemolytic activity (**Figure 2D**). The antibiotic susceptibility assay revealed that the strain E2 is sensitive to Tetracycline, Clindamycin, Penicillin, Cefalexin, Erythromycin, Chloramphenicol and Amoxicillin, yet showed antibiotics resistance to Vancomycin, Enrofloxacin, Kanamycin and Norfloxacin (**Table 1**). Thus, it was primarily considered as a safe probiotic for further *in vivo* experimental trials.

**Table 1.**
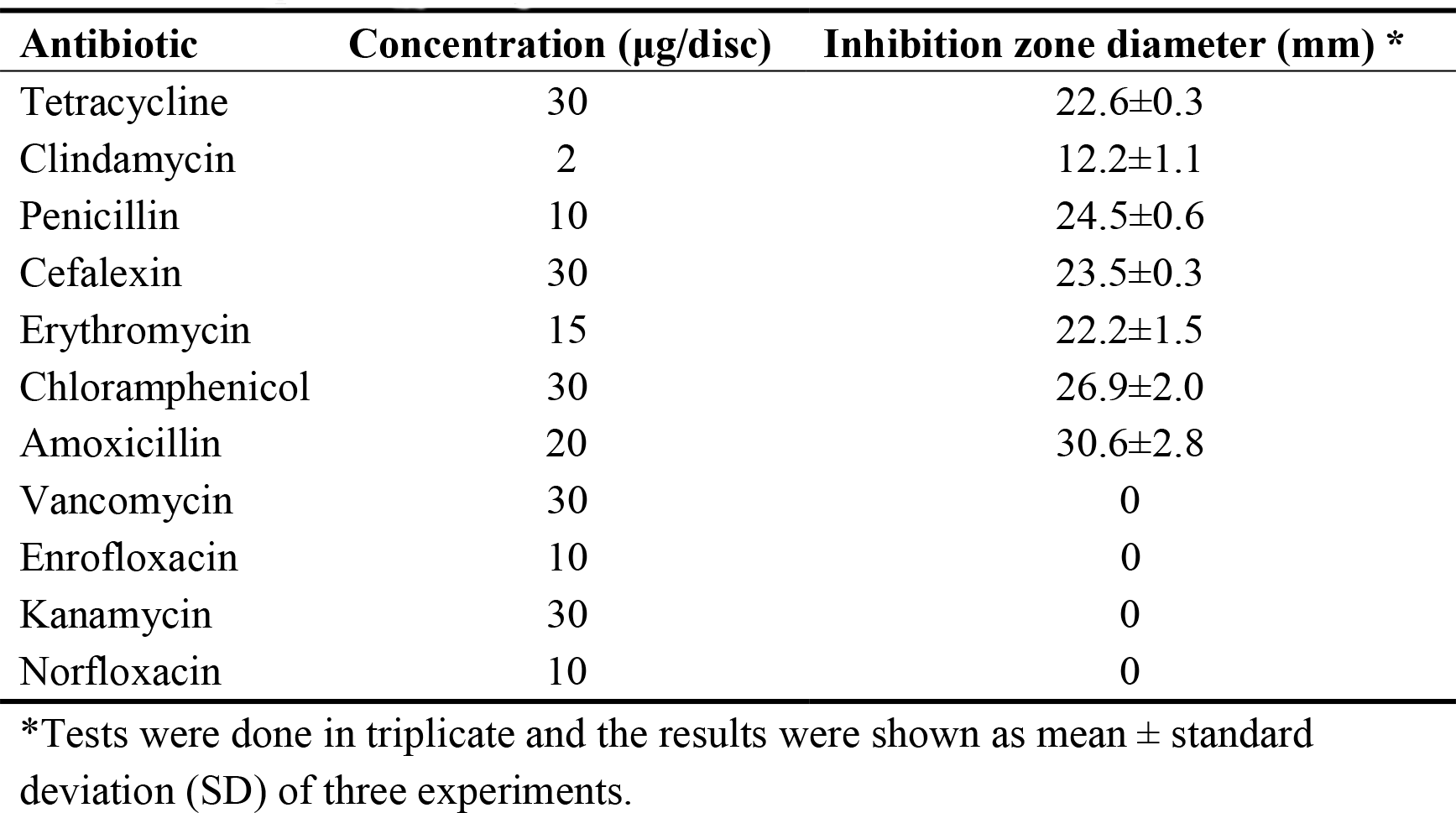
Susceptibility of *L. plantarum* strain E2 to commercial antibiotic discs.

### Effects of strain E2 on growth performance and survival of large yellow croaker

After 7 weeks of feeding, the E2 dietary supplementation significantly affected (*p* < 0.05) the growth rate of large yellow croakers (**Table 2**). The final body weight, final body length, weight gain rate and specific growth rate of fish in the E2 group were significantly higher than those in the control group (*p* < 0.05).

**Table 2.** Growth rate of large yellow croakers fed on the diet supplemented with *L. planturum* E2 for 7 weeks.

| Items | Control group | E2 group |
| --- | --- | --- |
| Initial body weight (g) | 3.15±0.07 | 3.20±0.07 |
| Final body weight (g) | 6.20±0.17 | 7.03±0.18* |
| Initial body length (cm) | 7.59±0.06 | 7.71±0.05 |
| Final body length (cm) | 8.53±0.08 | 9.58±0.09** |
| Weight gain rate (%) | 96.98±1.35 | 119.75±1.70*** |
| Specific growth rate (% day<sup>-1</sup>) | 1.61±0.01 | 1.88±0.02*** |

Data are expressed as mean ± S.E.M from a triplicate set of 200 fish (n = 3).
\* $p < 0.05$ , \*\* $p < 0.01$ , \*\*\* $p < 0.001$ .

### Effects of strain E2 on gut health and digestive ability of large yellow croaker

To explore the effects of strain E2 on gut health of large yellow croaker, intestinal structure of fish in the control group and E2 group was analyzed by histology method. As shown in Figure 3A, compared to the control group, fish in the E2 group exhibited a more complete and healthier gut structure, as evidenced by higher villus height and reduced pigment deposition in the lamina propria. Moreover, the strain E2 also decreased the edema and inflammatory cell infiltration in the gut lamina propria (**Figure 3A**).

**Figure 3.**
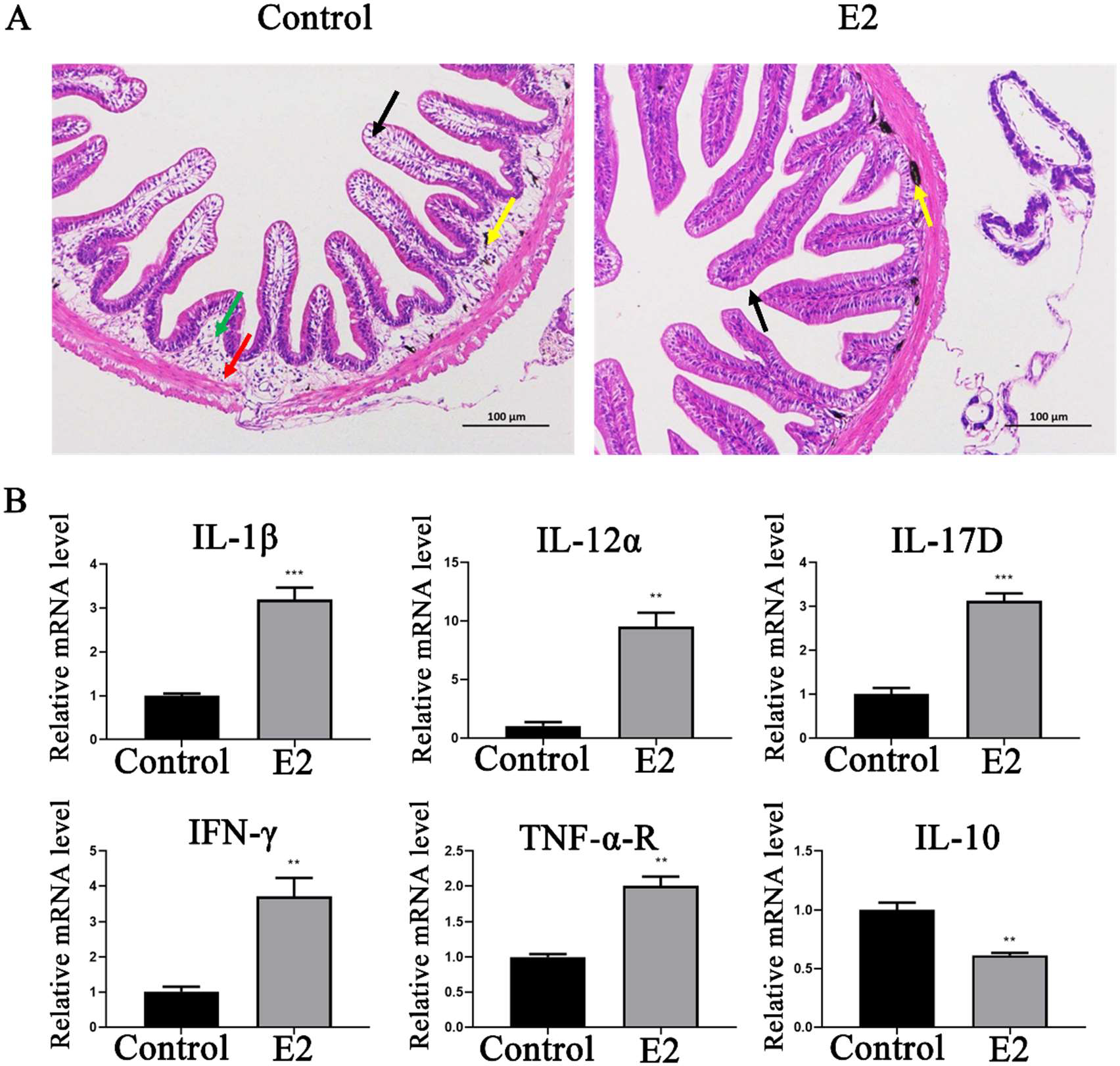
Effects of *L. plantarum* strain E2 on intestinal structure and immune-related gene expression in large yellow croaker. (A) Intestinal morphology observation. Views under 20 × objective lens (HE staining; Scale bar = 100 μm). Black row, villus height; yellow row, pigment deposition; green row, edema in gut lamina propria; red row, inflammatory cell infiltration. (B) The expression changes of the immune-related genes in the intestinal tissue of fish in the E2 group. Data represent the means ±SEM (n = 3). **, *p* < 0.01; ***, *p* < 0.001.

The strain E2 dietary supplementation could affect the gut digestive ability of fish. The intestinal α-amylase and trypsin activities of fish in the E2 group were significantly higher than those in the control group (*p* < 0.001; *p* < 0.05) (**Table 3**), while the lipase activity in E2 group was also higher than that in control group despite no statistical difference observed (*p* > 0.05) (**Table 3**).

**Table 3.** The digestive enzyme activity of large yellow croakers fed with the diet supplemented with *L. planturum* E2.

| Items | Control group | E2 group |
| --- | --- | --- |
| α-Amylase (U/mgprot) | 0.611±0.024 | 0.789±0.013*** |
| Trypsin (U/mgprot) | 474.468±104.199 | 1254.915±148.221* |
| Lipase (U/mgprot) | 0.272±0.062 | 0.537±0.092 |

Data are presented as means ± S.E.M (n = 3). \* $p$ < 0.05, \*\* $p$ < 0.01, \*\*\* $p$ < 0.001.

Intestinal tissue was selected to investigate the expression of six immune-related genes (IL-1β, IL-12α, IL-17D, IFN-γ, TNF-α-R and IL-10) using the β-actin as the endogenous reference (Mao et al., 2018). The strain E2 supplementation significantly decreased the mRNA expression of IL-10 (*p* < 0.01) compared with the control group and the mRNA levels of genes related to immune response were increased significantly (*p*<0.01). It is worth noting that feeding with E2 resulted in a high expression of IL-12 in the intestine and the IFN-γ regulated by it was correspondingly upregulated significantly. Studies have shown that bacteria have a modulatory effect on IL-12 (Kamada et al., 2013), so it is speculated that it may due to the addition of *L. plantarum* E2 to stimulated IL-12 expression in the intestine, making the E2 group showing an immune enhancing effect. As shown in Figure 3B, strain E2 supplementation significantly increased the expression level of IL-1β, IL-12α, IL-17D, IFN-γ and TNF- α-R.

### Effects of E2 on gut microbiota of large yellow croaker

After the quality control and trimming, a total of 523,773 high quality clean reads from fish in the control group and E2 group were obtained, resulting in the identification of 94 OTUs. The OTUs were assigned to 9 phyla and 38 genera. At the phyla level, Venn diagram showed that two groups shared 30 OTUs. Control group and E2 group possessed 35 and 29 unique OTUs respectively (**Figure 4A**). At the genus level, the two groups shared 19 genera, while control group and E2 group possessed 10 and 9 unique OTUs respectively (**Figure 4B**).

**Figure 4.**
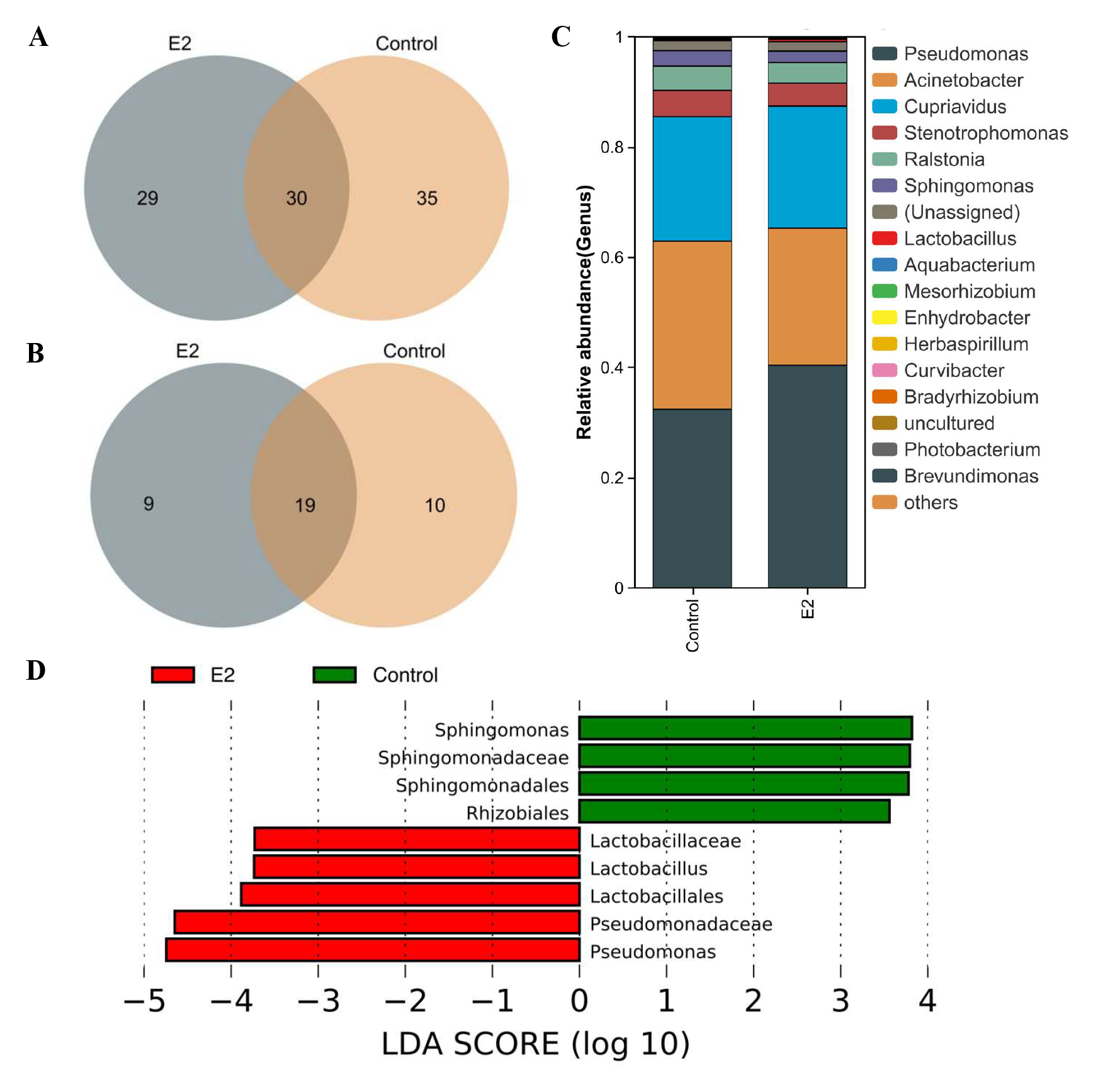
Effects of *L. plantarum* E2 on gut microbiota of large yellow croaker. **(A)** Venn diagram at phylum taxonomic levels. **(B)** Venn diagram at genus taxonomic levels. **(C)** Classification of reads at genus taxonomic levels. Only top 10 most abundant (based on relative abundance) bacterial genera were shown. Other genera were all assigned as ‘Others’. **(D)** LEfSe analysis identified the most differentially abundant taxons between control and *L. plantarum* E2 groups (n = 3/group). Histogram of linear discriminant analysis (LDA) scores for differentially abundant taxons. Cladogram was calculated by LEfSe, and displayed based on the effect size.

The microbial community composition analysis revealed the differences in gut microbiota between two groups. At the genus level, *Pseudomonas*, *Acinetobacter* and *Cupriavidus* were most predominant bacteria in both E2 group and control group. However, *Lactobacillus* and *Mesorhizobium* had higher abundance only in the E2 group, while *Enhydrobacter* and *Curvibacter* were only in the top 10 predominant bacteria of the control group (**Figure 4C and Supplementary Table 6**).

To further explore the influence of E2 on gut microbiota of large yellow croaker, LEFSe analysis was used to identify the difference in microbial community composition between two groups. The results showed that the great changes have taken place in taxonomic distribution of gut microbiota after E2 diet. E2 diet significantly decreased the relative abundance of order *Sphingomonadales*, family *Sphingomonadaceae*, genus *Sphingomonas* and order *Rhizobiales* (*p* < 0.05) (**Figure 4D**). Some bacteria in the order *Lactobacillales*, family *Lactobacillaceae,* genus *Lactobacillus* and order *Pseudomonadales*, family *Pseudomonadaceae*, genus *Pseudomonas* were increased after E2 diet feeding (*p* < 0.05).

### Effects of strain E2 on the survival of large yellow croaker after challenge with *Pseudomonas plecoglossicida* PQLYC4

The survival of large yellow croakers was recorded within 6 days after the artificial challenge with *P. plecoglossicida* PQLYC4, and the survival curve was drawn. As shown in Figure 5, the strain E2 dietary supplementation obviously improved the resistance of large yellow croaker against *P. plecoglossicida* PQLYC4 (*P* < 0.05).

**Figure 5.**
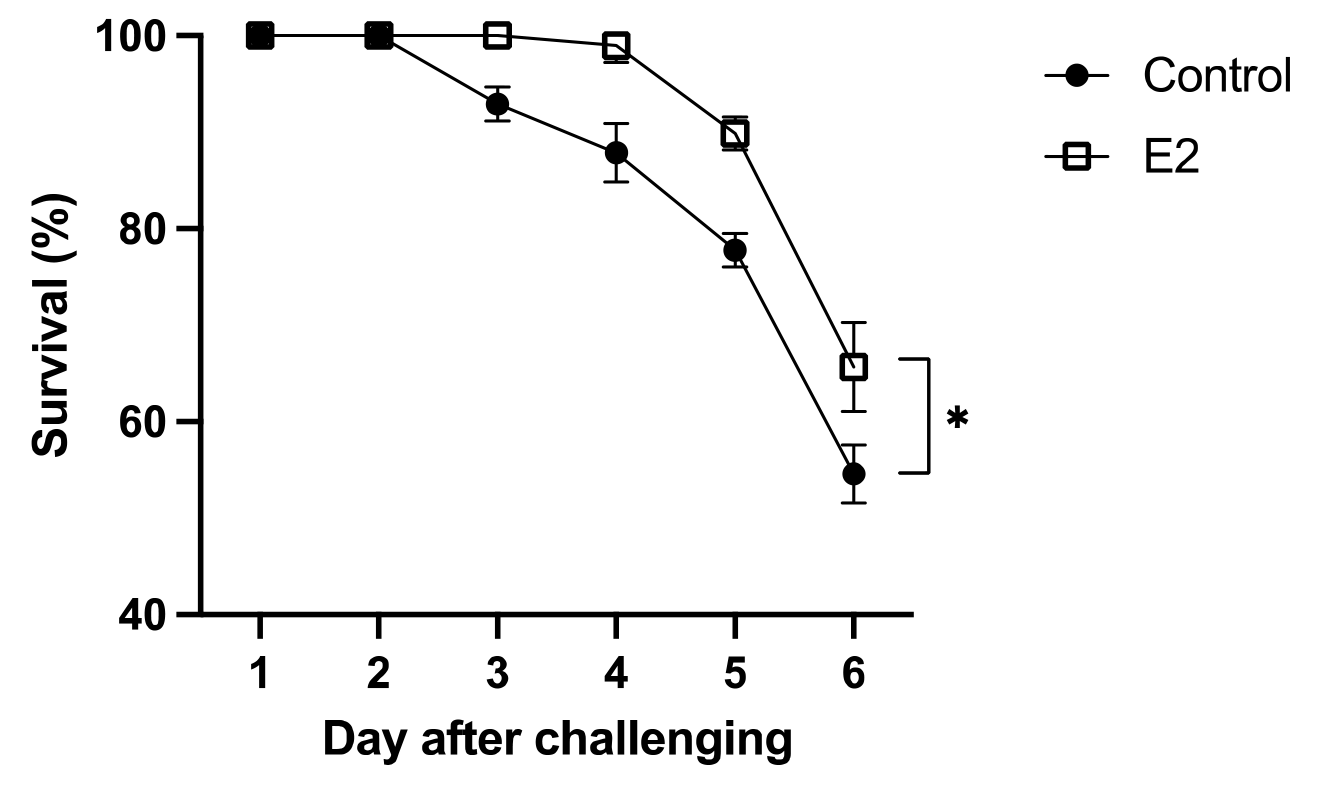
Survival rate of large yellow croakers after challenge with *P. plecoglossicida* PQLYC4. Data represent the means ± SEM (n = 3). *, *p* < 0.05

## Discussion

Bacterial diseases have caused the substantial economic losses to aquaculture, which has become one of the largest bottlenecks that restrict the sustainable development of aquaculture industry (Lin et al., 2022; Rodger, 2016). With the increasingly strict supervision of antibiotics or other chemical agents, probiotics are considered more and more important for controlling bacterial diseases in aquaculture (Chauhan and Singh, 2019; W. Zhang et al., 2020). In this study, a bacterial strain E2 isolated from the intestinal tract of cultured large yellow croaker showed significant antibacterial activity. After taxonomy identification, the strain E2 was identified as *L. plantarum. L. plantarum* is wildly distributed in nature, including the intestinal tract of insects, mammals, humans, and fish (Iorizzo et al., 2021, 2020; Nordström et al., 2021; Siezen et al., 2010). Several strains of *L. plantarum,* such as strain LB411, L34b-2, and 73V, have been reported to show multiple antibacterial activities against *Escherichia coli, Listeria innocua, Proteus mirabilis, Staphylococcus aureus, Pseudomonas aeruginosa, Citrobacter freundii,* and *Aeromonas salmonicida* (Alonso et al., 2019; Iorizzo et al., 2022; Meidong et al., 2021). *L. plantarum* E2 isolated here could suppress five fish pathogenic bacteria, including *P. plecoglossicida, V. campbellii, V. harveyi, V. alginolyticus,* and *A. hydrophila,* showing a broad spectrum of antibacterial activity. Due to being isolated from the intestinal tract of large yellow croaker, *L. plantarum* E2 also showed excellent biological safety and tolerance to acidic environments and bile salts. These characteristics of *L. plantarum* E2 were similar to the other strains of *L. plantarum* isolated from the intestinal tracts of *Salmo macrostigma* (Iorizzo et al., 2022).

Probiotics can promote the growth of fish. Here, *L. plantarum* E2 showed satisfactory growth improvement in large yellow croaker. After 7 weeks of feeding, the E2 supplementation of dietary significantly affected the final body weight, final body length, weight gain rate and specific growth rate of fish. The promotion on growth performance of fish were also observed in *L. plantarum* AH78 supplemented dietary which could significantly increase specific growth rates, feed conversion ratio, protein efficiency ratio, and productive protein value of Nile tilapia (Hamdan et al., 2016). *L. plantarum* CR1T5 also showed a significant increase in specific growth rate, weight gain and final weight of Nile tilapia (Van Doan et al., 2019). The similar growth improvement of other *L. plantarum* strains were found in Rohu (*Labeo rohita*), European sea bass (*Dicentrarchus Labrax*), juvenile turbot (*Scophthalmus maximus* L.) and grouper (*Epinephelus coioides*) (Giri et al., 2013; Piccolo et al., 2015; Son et al., 2009; L. Wang et al., 2016).

The intestinal mucosal barrier plays a critical role in the maintenance of fish health (Hongling Zhang et al., 2020). In large yellow croaker, *L. plantarum* E2 supplementation effectively improved villus height and decreased pigment deposition, edema and inflammatory cell infiltration in the gut lamina propria. These results were consisted with the previous studies on *L. plantarum* JCM1149, AH 78, 08.923 and WCFS1 in Nile tilapia and zebrafish (Hamdan et al., 2016; Li et al., 2022; Liu et al., 2016; Y. Wang et al., 2016). However, the mechanism of maintaining intestinal structural integrity by probiotics remains unclear. Current opinion suggested that the integrity of intestinal structure of farmed fish closely related with balance of nutrient absorption and immunity (Hongling Zhang et al., 2020). In this study, *L. plantarum* E2 significantly increased intestinal α-amylase, trypsin and lipase activities to improve digestion of large yellow croaker. Moreover, strain E2 also significantly suppressed the expression of anti-inflammatory cytokine IL-10 and increased the expression of pro-inflammatory cytokine IL-1β, IL-12α, IL-17D, IFN-γ and TNF-α-R. These results were same as the previous study of the effects of *L. plantarum* AH 78 supplementation in Nile tilapia (Hamdan et al., 2016). The strain E2 increased the expression of immune genes in large yellow croaker, allowing the organs to respond more rapidly to the pathogens, and the increase in the height of intestinal villus and the improvement in the morphology of the intestine may be attributed to the probiotics promoting the maturation of epithelial cells and improving the enhancement of nutrient absorption.

Gut microbiota can regulate nutrition, metabolism, immunity, and health of fish (Clements et al., 2014; Ray et al., 2012). A recent study indicated that modulation of intestine microbiota in farmed fish towards potentially more beneficial microbial community could be achieved through probiotics supplementation (Yin et al., 2021). In this study, the addition of *L. plantarum* E2 significantly increased bacteria of genus *Lactobacillus* and *Pseudomonas*. Previous studies have revealed that some strains of *Pseudomonas* were probiotics. *P. synxantha* and *P. aeruginosa* were reported to benefit on the growth, survival and immune parameters of juvenile western king prawns (Hai et al., 2009). In addition, *L. plantarum* E2 also significantly decreased the relative abundance of genus *Sphingomonas.* Evidence have shown that several *Sphingomonas* species are known opportunistic pathogens (Hyun et al., 2021; Pękala-Safińska, 2018). Generally, the ratio of probiotics to pathogens in the intestinal tract will affect the food digestion and immunity of host (Merrifield et al., 2010). Therefore, the *L. plantarum* E2 induced alteration in gut microbiota may promote gut health in large yellow croaker.

In conclusion, our findings demonstrate that *L. plantarum* E2 isolated from the intestinal tract of large yellow croaker has significant growth promotion and disease resistance of large yellow croaker as dietary supplements. The beneficiary effects of *L. plantarum* E2 might achieved though the maintenance of intestinal mucosal barrier, promotion of intestinal digestion and inhibiting inflammatory response. Furthermore, *L. plantarum* E2 also altered gut microbiota to promote gut health in large yellow croaker. These results indicated that *L. plantarum* E2 has potential application as a probiotic in aquaculture.

## Declaration of Competing Interest

The authors declare that they have no conflict of interest.

## Author contributions

Ruizhe Liu performed most of the experiments, analyzed the data, and wrote the manuscript. San Wang, Dongliang Huang, Yulu Huang help with experimental operations. Tianliang He and Xinhua Chen designed the research and revised the manuscript. All authors contributed to the article and approved the submitted version.

## Acknowledgement

This work was supported by National Key Research and Development Program of China Grants 2022YFD2401001, National Natural Science Foundation of China Grant U1905204, China Agriculture Research System of MOF and MARA Grant CARS-47, and Fujian Science and Technology Department Grants 2021N5008.

## Notes

### Competing Interest Statement

The authors have declared no competing interest.

